# Degraded tactile coding in the Cntnap2 mouse model of autism

**DOI:** 10.1101/2023.09.29.560240

**Authors:** Han Chin Wang, Daniel E. Feldman

## Abstract

Atypical sensory processing in autism involves altered neural circuit function and neural coding in sensory cortex, but the nature of coding disruption is poorly understood. We characterized neural coding in L2/3 of whisker somatosensory cortex (S1) of *Cntnap2^-/-^* mice, an autism model with pronounced hypofunction of parvalbumin (PV) inhibitory circuits. We tested for both excess spiking, which is often hypothesized in autism models with reduced inhibition, and alterations in somatotopic coding, using c-fos immunostaining and 2-photon calcium imaging in awake mice. In *Cntnap2^-/-^* mice, c- fos-(+) neuron density was elevated in L2/3 of S1 under spontaneous activity conditions, but comparable to control mice after whisker stimulation, suggesting that sensory-evoked spiking was relatively normal. 2-photon GCaMP8m imaging in L2/3 pyramidal cells revealed no increase in whisker-evoked response magnitude, but instead showed multiple signs of degraded somatotopic coding. These included broadening of whisker tuning curves, blurring of the whisker map, and blunting of the point representation of each whisker. These altered properties were more pronounced in noisy than sparse sensory conditions. Tuning instability, assessed over 2-3 weeks of longitudinal imaging, was also significantly increased in *Cntnap2^-/-^* mice. Thus, *Cntnap2^-/-^* mice show no excess spiking, but a degraded and unstable tactile code in S1.

## Introduction

How neural circuit dysfunction leads to the sensory and cognitive phenotypes of autism spectrum disorder (ASD) remains elusive. Because many ASD risk gene mutations reduce parvalbumin (PV) interneuron function and increase excitation-inhibition ratio (E-I ratio) in cerebral cortex, a dominant circuit-level model has been that cortical circuits exhibit hyperexcitability and excess spiking in ASD. Such excess spikes could increase noise in neural coding, thus impairing information processing (Rubenstein and Merzenich, 2003; Sohal and Rubenstein, 2019). Alternatively, PV hypofunction or other ASD-related circuit changes could drive other forms of neural coding impairment without any excess spiking—for example, broadening neural tuning, degrading cortical maps, disrupting cortical rhythms or precise spike timing, or degrading the structure of population codes. Distinguishing ’degraded coding’ from ’hyperexcitability’ models of circuit dysfunction in ASD is critical for developing effective therapeutic approaches to autism.

Sensory cortex is a useful site to investigate circuit dysfunction in autism, because atypical sensory processing occurs in up to 90% of individuals with ASD (Levy et al., 2009). Across different transgenic mouse models of ASD, PV hypofunction is common in sensory cortex, but excess spiking is rare, and instead various forms of degraded neural coding have been observed in many models (Monday et al., 2023). Excess spiking occurs in *Shank3b^-/-^* mice, and in some sensory cortical areas in *Fmr1^-/y^* mice (Chen et al., 2020; Gonçalves et al., 2013; Rotschafer and Razak, 2014; Zhang et al., 2014). But spiking activity is normal or reduced in sensory cortex of many other ASD models, including *MeCP2^-/-^*, *Syngap1^+/-^*, *16p11.2* deletion, and other primary sensory cortical areas in *Fmr1^-/y^*, raising the question of what may underlie abnormal sensory detection or discrimination behavior in these models (Antoine et al., 2019; Banerjee et al., 2016; Goel et al., 2018; Juczewski et al., 2016; Michaelson et al., 2018). The goal of the current study is to characterize neural coding phenotypes in primary somatosensory cortex (S1) in the *Cntnap2^-/-^* mouse model of autism. We focused on S1 because tactile abnormalities are common in autism and may contribute to development of social deficits (Orefice et al., 2019; Robertson and Baron-Cohen, 2017; Schaffler et al., 2019), and *Cntnap2* is expressed in primary sensory organs and brain areas engaged in sensory processing (Gordon et al., 2016).

The *CNTNAP2* gene encodes Contactin-associated protein 2 (CASPR2), a cell adhesion molecule that localizes at the axon initial segment and nodes of Ranvier, and clusters Kv1 channels to regulate spiking excitability (Bonetto et al., 2019). *CNTNAP2* is an autism risk gene. Loss-of-function mutation of *CNTNAP2* causes Pitt-Hopkins-Like Syndrome 1 (PTHSL1, OMIM# 610042), which includes autistic symptoms, and single nucleotide polymorphisms (SNPs) in *CNTNAP2* gene are associated with increased susceptibility of autism (Alarcón et al., 2008; Arking et al., 2008; Bakkaloglu et al., 2008). *Cntnap2^-/-^* mice (Peñagarikano et al., 2011; Poliak et al., 2003) show social, communication and repetitive motion behavioral phenotypes (Brunner et al., 2015; Peñagarikano et al., 2011), as well as atypical tactile, auditory, olfactory and visual processing (Balasco et al., 2022; Dawes et al., 2018; Deemyad et al., 2022; Del Rosario et al., 2021; Gordon et al., 2016; Scott et al., 2019; Truong et al., 2015).

*Cntnap2^-/-^* mice show weakened synaptic inhibition and excitation in S1 and other cortical areas (Antoine et al., 2019; Bridi et al., 2017; Jurgensen and Castillo, 2015; Lazaro et al., 2019; Paterno et al., 2021; Sacai et al., 2020), and strong evidence for PV hypofunction (Antoine et al., 2019; Deemyad et al., 2022; Jurgensen and Castillo, 2015; Lauber et al., 2018; Vogt et al., 2018). In layer (L) 2/3 of S1, whisker-evoked spiking of PV cells and PV-mediated feedforward inhibition are reduced (Antoine et al., 2019). But whether *Cntnap2^-/-^* mice exhibit excess spiking in sensory cortex is unclear: Prior studies of single-unit spiking reported normal spiking in S1 (Antoine et al., 2019), weaker-than-normal spiking in V1 (Del

Rosario et al., 2021), but excess spiking in S1 and A1 inferred from c-fos staining and multi-unit spike recording (Balasco et al., 2022; Scott et al., 2022). We sought to resolve whether S1 exhibits excess spiking in *Cntnap2^-/-^* mice, and if not, to test for other forms of neural coding abnormalities that may occur.

Detection of coding abnormalities in S1 is aided by the detailed understanding of whisker tuning and whisker map organization in wild type mice. S1 neurons have narrow somatotopic tuning for one or a few neighboring whiskers on the face. L4 barrels mark anatomical columns, one for each whisker in a precise anatomical somatotopic map (Woolsey and Loos, 1970). Within L4, nearly all neurons are tuned for their columnar whisker. In L2/3, pyramidal (PYR) neurons tuned to different whiskers are intermixed in each column, but the most common tuning is for the columnar whisker, resulting in a L2/3 whisker map with correct average topography, but high local scatter (Clancy et al., 2015; LeMessurier et al., 2019; Wang et al., 2022). As a result, the set of neurons tuned for a given whisker is distributed across several columns, centered on the whisker’s anatomical column. Interestingly, the somatotopic tuning of many L2/3 PYR neurons is markedly unstable over days and weeks, an example of representational drift (Wang et al., 2022). Such tuning drift is found in several primary sensory cortical areas as well as high-order association cortices, and may represent a challenge for maintaining stable cortical population codes (Deitch et al., 2021; Schoonover et al., 2021; Ziv et al., 2013).

We characterized sensory coding in S1 of *Cntnap2^-/-^*and *Cntnap2^+/+^* mice, using 2-photon calcium imaging in awake mice that received calibrated whisker stimuli as they performed a sensory task. We also used c-fos immunostaining to assess gross changes in neural activity in S1. Results showed that *Cntnap2^-/-^* mice do not exhibit excess sensory-evoked activity in L2/3 of S1. Instead, they show several forms of degraded sensory coding, including broadened single-neuron tuning, substantially degraded map topography, and elevated tuning instability.

## Results

### Whisker-evoked activity in L2/3 of S1 measured by c-fos expression

We assayed for gross alterations in S1 cortical activity in *Cntnap2^-/-^* mice by expression of the immediate early gene *c-fos* (Dragunow and Faull, 1989; Filipkowski et al., 2000) in L2/3. Mice were lightly anesthetized and head-fixed, and whiskers on the right side of the face in the A, C and E rows, plus associated beta and delta whiskers, were inserted into a piezoelectric actuator array. Calibrated deflections were applied to these whiskers for 30 min (0.5-sec deflection trains, delivered every 5 sec). After 60-90 min recovery, mice were euthanized, and flattened sections were made from S1, parallel to layer 4, from both the contralateral hemisphere (whisker stimulated) and the ipsilateral hemisphere (treated as an unstimulated control). Sections were processed for c-fos immunofluorescence and co-stained with streptavidin to reveal L4 barrels (Sumser et al., 2017).

c-fos positive cells were identified in L2/3, and their positions were marked relative to barrel column boundaries, identified from the L4 barrels (Fig. 1a-b). In the contralateral (stimulated) hemisphere, the density of c-fos positive cells was higher in L2/3 of cortical columns corresponding to deflected A, C and E whiskers, relative to columns for undeflected B and D whiskers, in both *Cntnap2^+/+^* (control) and *Cntnap2^-/-^* mice (Fig. 1b-c) (control mice: p<0.001,unbalanced two-way ANOVA, n=9 columns in 3 mice; Cntnap2^-/-^: p<0.001,unbalanced two-way ANOVA ,n=9 columns in 3 mice ). In the ipsilateral (unstimulated) hemisphere, c-fos cell density was equal in A-C-E vs. B-D whisker columns (Fig. 1c) (control mice: p=0.383; Cntnap2^-/-^: p=0.522). Thus, 30 min of whisker stimulation effectively activates L2/3 neurons, as assayed by c-fos expression.

**Figure 1.**
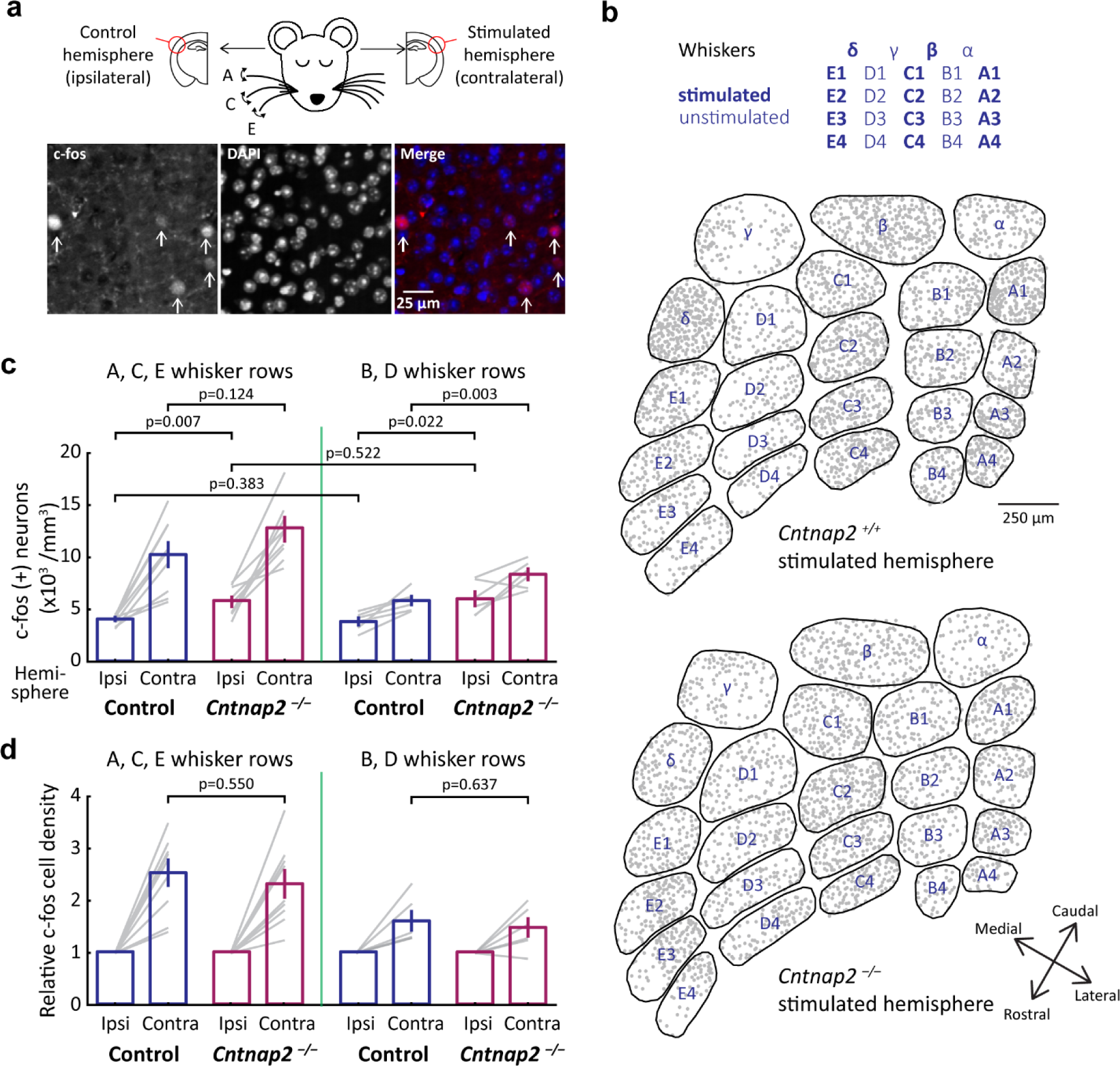
Spontaneous and whisker-evoked c-fos expression in L2/3 of S1 in *Cntnap2^+/+^* and *Cntnap2^-/-^*mice **(a).** Top: Design of c-fos immunolabeling experiment. Bottom: Examples of cells co-labeled with c-fos and nuclear stain (DAPI). **(b).** Locations of all c-fos-positive L2/3 neurons relative to L4 barrel boundaries in the contralateral (stimulated) hemisphere from one control and one Cntnap2^-/-^ mouse. Data were compiled across 250 µm in L2/3. **(c).** Density of c-fos-positive cells in L2/3 of S1, comparing stimulated and unstimulated hemisphere, across all mice. Each gray line is one column (A, B, C, D, or E) in one mouse. p-values are for genotype factor in unbalanced two-way ANOVA. **(d).** Same data as (c), normalized to the unstimulated hemisphere of each mouse. p-values are for genotype factor in unbalanced two-way ANOVA.

In the ipsilateral, unstimulated hemisphere, *Cntnap2^-/-^*mice showed increased density of c-fos-positive cells in both A-C-E and B-D whisker rows relative to *Cntnap2^+/+^* mice, suggesting higher spontaneous activity (Fig. 1c, ipsilateral; p-values shown in figure). In the contralateral, stimulated hemisphere, whisker stimulation evoked similar c-fos density in A-C-E columns of *Cntnap2^+/+^* and *Cntnap2^-/-^* mice, with only a modest, non-significant trend for increased density in *Cntnap2^-/-^* mice (Fig. 1c, left). Whisker stimulation also modestly increased c-fos cell density in the B and D rows of the stimulated hemisphere, consistent with salt-and-pepper somatotopy in L2/3 of S1 (Clancy et al., 2015; LeMessurier et al., 2019; Pluta et al., 2017; Wang et al., 2022). *Cntnap2^-/-^* mice showed a greater density of c-fos cells in B-D rows of the stimulated hemisphere than control mice, suggesting that the whisker map might become more spatially distributed or blurred in L2/3 of *Cntnap2^-/-^*mice (Fig. 1c, right).

We normalized c-fos cell density in each contralateral hemisphere to the ipsilateral, unstimulated hemisphere of the same mouse, in order to infer signal-to-noise ratio for whisker-evoked neural activation over spontaneous activity. *Cntnap2^+/+^* and *Cntnap2^-/-^*mice showed indistinguishable multiplicative increases in c-fos cell density after whisker stimulation (Fig. 1d), both for A-C-E and B-D columns, suggesting that the signal-to-noise ratio for sensory activation was not altered in *Cntnap2^-/-^* mice, even though the number of spontaneously active cells was greater. Together, these c-fos results suggests increased spontaneous activity and a more dispersed whisker map in S1 of *Cntnap2^-/-^* mice, but no significant increase in the magnitude of whisker-evoked sensory activation.

### Whisker discrimination task for 2-photon calcium imaging in S1 of awake mice

To characterize sensory coding more accurately in *Cntnap2^-/-^*mice, we performed Ca^2+^ imaging of whisker-evoked activity in L2/3 of S1 in awake, whisker-attentive mice. We trained 10 mice on a behavioral task that allows quantitative assessment of whisker-evoked responses, receptive fields, and maps by 2-photon imaging during task performance (Wang et al., 2022). Head-fixed mice had 9 whiskers inserted in a 3 x 3 piezoelectric actuator array. Each piezo stimulated one whisker with a brief train of whisker deflections (termed a whisker cue, 5 impulses at 100 ms inter-impulse interval). On each trial, one of 11 stimuli was presented: either one of the 9 single-whisker cue stimuli, a blank (no stimulus), or an all-whisker stimulus (Fig. 2a). Mice learned to lick to the all-whisker stimulus, which was rewarded (S+), but not to single-whisker cues or blanks, which were not rewarded (S–). Once mice are trained, this task design allows single-whisker cue stimuli (S–) to be used to map whisker responses and receptive fields without lick contamination or the need for a delay period.

**Figure 2.**
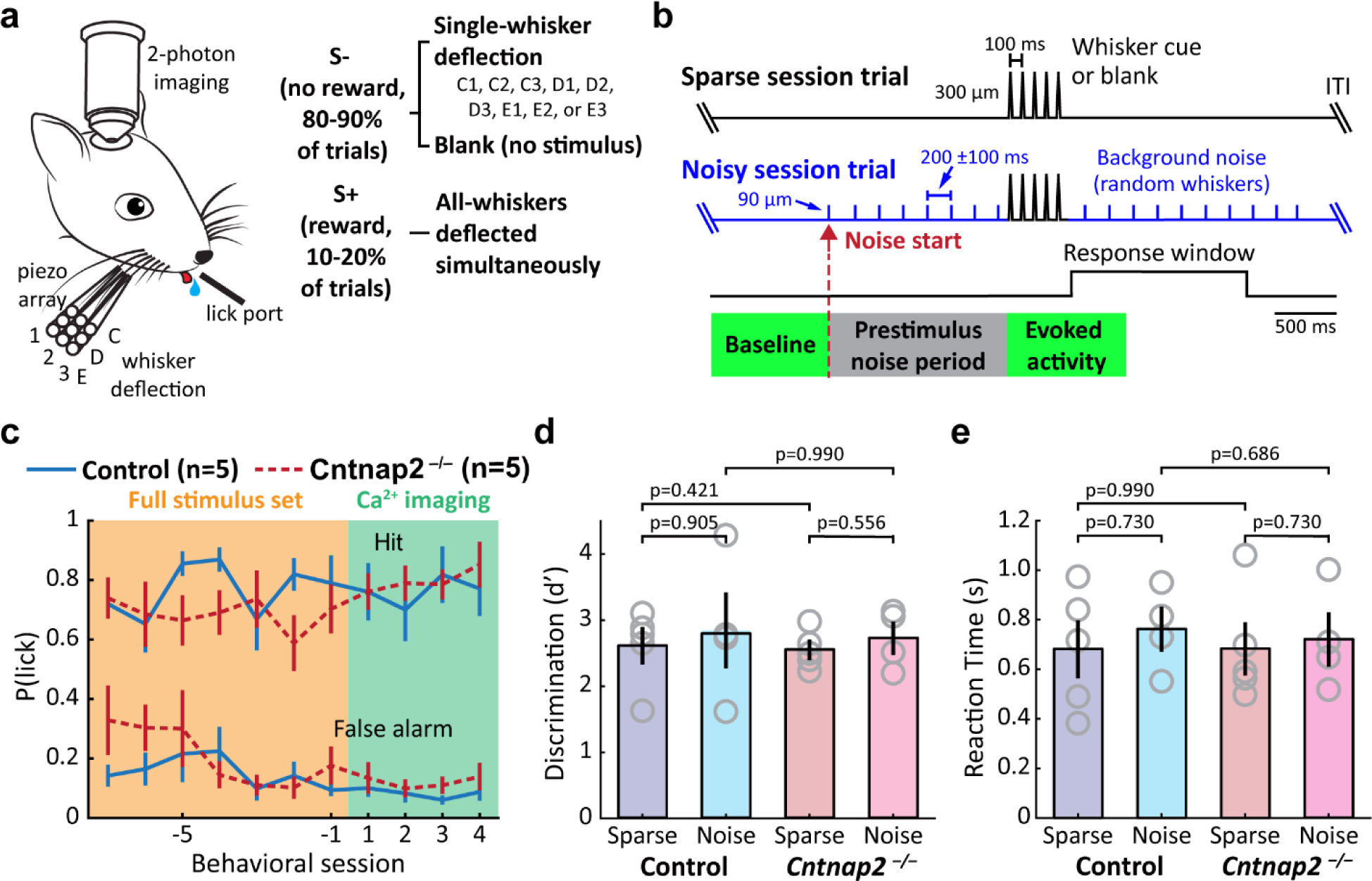
Behavior task for receptive field mapping in awake mice**(a).** Experiment design and setup, showing stimuli and reward assignment for the task. **(b).** Trial structure for sparse and noisy sessions. **(c).** Behavioral performance during training, when all S+ and S– stimuli are present (orange, showing the last 7 sessions prior to imaging), and once mice have achieved expert performance and imaging begins (green, showing the first 4 imaging sessions). n: animals. **(d).** Behavioral discrimination performance (d-prime) during imaging sessions, by genotype and noise condition. Statistics: Rank-sum. **(e).** Reaction time for hit trials, by genotype and noise condition. Statistics: Rank-sum. All error bars are SEM.

We hypothesized that excess spiking or degraded sensory coding may emerge in noisy sensory conditions, compared to sparse, quieter sensory conditions. To test this, we interleaved two types of behavioral sessions (Fig. 2b). In sparse stimulus sessions, each trial contained only one whisker cue, amounting to one cue stimulus every 6.5 ± 1 seconds. In noisy stimulus sessions, small single-whisker deflections (30% of whisker cue amplitude) were applied on random, interleaved whiskers every 200 ± 100 ms throughout the trial, starting 1.5 seconds before whisker cue delivery. The period of noise stimulation prior to whisker cue delivery was called the “prestimulus noise period”. The goal of this design was to raise sensory background noise, thus complicating encoding and making detection more difficult. Mice were trained in the sparse condition only, after which sparse and noise sessions were interleaved (only one condition was tested per day).

Mice reached expert performance in the sparse condition, defined as hit rate > 70% and false alarm rate < 25%, in 10.1 ± 0.5 days after introduction of all S– stimuli (n = 10 mice). There was no difference in this training duration between genotypes (*Cntnap2^+/+^*: 10 ± 0.6 days; *Cntnap2^-/-^*: 10.2 ± 0.9 days, n = 5 mice each) (Fig. 2c). There was also no difference in detection performance, measured by d-prime, or in reaction time, between expert *Cntnap2^+/+^* and *Cntnap2^-/-^* mice, or between sparse and noisy conditions (Fig. 2d-e).

### L2/3 PYR neurons show more spontaneous activity and slightly broader tuning, but no evidence of sensory-evoked hypersensitivity

We measured sensory responses in L2/3 pyramidal (PYR) cells by virally expressing GCaMP8m (Zhang et al., 2023), and imaging in L2/3 during the task. Whisker-evoked responses and receptive fields were measured from whisker cue stimuli on the single-whisker S– trials. Trials with licks (i.e., false alarm trials) were excluded from analysis to avoid lick-related neural activity and motion artefacts. Imaged neurons were localized post-hoc relative to anatomical column boundaries by reconstructing each imaging field relative to barrels in L4 (Golshani et al., 2009; Wang et al., 2022). Whisker stimuli evoked strong ΔF/F responses from L2/3 PYR cells, allowing us to assess whisker-evoked response magnitude and whisker tuning of each cell (Fig. 3a-b).

**Figure 3.**
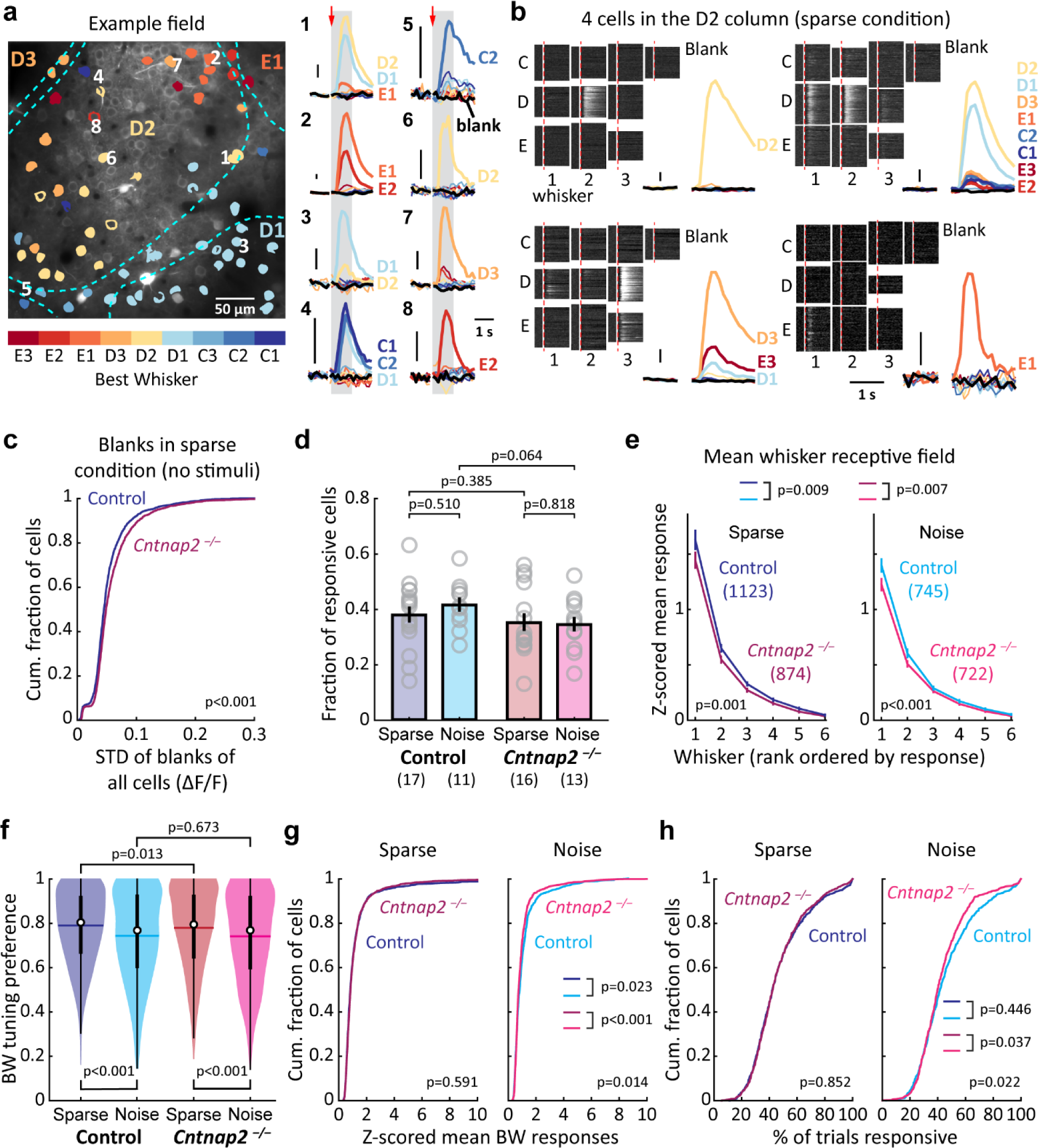
Whisker responsiveness and tuning of L2/3 PYR neurons in *Cntnap2^+/+^* and *Cntnap2^-/-^* mice **(a).** Example imaging field showing GCaMP8m-expressing PYR cells color-coded for their BW. Dashed lines, barrel boundaries from L4. Right: mean whisker-evoked ΔF/F traces for 8 cells. Thick traces and whisker labels show significant whisker responses; thin traces are non-significant responses; black traces are blanks. Arrows showed whisker deflection onset. Gray, response analysis window. Scale bar: 0.1 ΔF/F. **(b).** Whisker tuning and trial-to-trial reliability for 4 example cells imaged in the D2 column. Single- trial ΔF/F traces for each single-whisker trial and blank trial are shown, with the mean whisker-evoked ΔF/F traces for that cell. Dash, stimulus onset. ΔF/F traces are normalized to maximum for that cell. Scale bar: 0.1 ΔF/F. **(c).** Standard deviation (STD) of responses in blank trials in sparse condition, as a measure of spontaneous activity. Statistics: KS. **(d).** Fraction of whisker-responsive neurons in each imaging field, by genotype and noise condition. Each circle is one imaging field. statistics: rank-sum. **(e).** Mean rank-ordered whisker tuning curves across all whisker-responsive neurons. Only cells whose BW and at least 5 adjacent whiskers were in the piezo array were included. Responses are z-scored to activity in blank trials. Error bars: SEM. p-values are for genotype or noise level factor in unbalanced two-way ANOVA. **(f).** Distribution of mean BW tuning preference of responsive neurons. Circles are medians, horizontal lines are means, thick vertical lines are interquartile ranges, and thin vertical line is 1.5X interquartile ranges. Statistics: rank-sum. **(g).** The cumulative distribution of z-scored BW responses of responsive neurons. Statistics: KS. **(h).** Cumulative fraction of individual trials with significant whisker response. Statistics: KS.

We analyzed 4334 PYR cells from *Cntnap2^+/+^* mice (2751 in sparse sessions, n = 5 mice and 1583 from noise sessions, n = 4 mice) and 3951 cells from *Cntnap2^-/-^* animals (2136 in sparse sessions, n = 5 mice and 1815 in noise sessions, n = 4 mice). Average ΔF/F in the prestimulus noise period was higher in noise sessions than in sparse sessions, suggesting that noise stimuli were effective in driving S1 activity (Fig. 3- Figure Supplement 1a). This background whisker noise could potentially interfere with neural coding of whisker information, even though behavioral detection of cue stimuli was equivalent in sparse and noise sessions (Fig. 2d-e).

To assess spontaneous activity, we calculated the standard deviation of ΔF/F for each cell within blank trials in the sparse condition, when no stimuli are present. *Cntnap2^-/-^* mice showed modestly but significantly higher values of this metric than control mice, indicating higher spontaneous activity (Fig. 3c), consistent with the c-fos results. Next, we examined responses to single-whisker cue stimuli.

Overall, ∼30-40% of PYR neurons were significantly whisker responsive, as expected from known sparse coding in S1 (Crochet et al., 2011; O’Connor et al., 2010; Wang et al., 2022). The fraction of responsive cells was not significantly different between *Cntnap2^+/+^* and *Cntnap2^-/-^*genotypes, or between sparse and noise conditions (*Cntnap2^+/+^*sparse: 37.9%; *Cntnap2^+/+^* noisy: 41.6%; *Cntnap2^-/-^*sparse: 35.2%; *Cntnap2^-/-^* noisy: 34.5%) (Fig. 3d).

Whisker somatotopic receptive fields were generally narrow, such that for each cell, only 1-3 whiskers drove ΔF/F responses that were significantly greater than on blank trials (example: Fig. 3a-b). Most cells had a single best whisker (BW) which elicited a statistically stronger response than any other whisker (*Cntnap2^+/+^* sparse: 57.6% of cells; *Cntnap2^+/+^*noisy: 53.0%; *Cntnap2^-/-^* sparse: 60.2%; *Cntnap2^-/-^*noisy: 55.3%; no differences between genotype or noise condition). The remaining cells had several (usually 2- 3) statistically co-equal best whiskers. Mean rank-ordered whisker tuning curves revealed that *Cntnap2^-/-^* mice had a slightly blunted tuning peak but normal flanks (Fig. 3e), causing a modest broadening of whisker tuning around the BW (assessed by BW preference metric, Fig. 3f). This was observed in both sparse and noisy conditions (Fig. 3f).

To test for sensory hypersensitivity, we identified the absolute BW for each cell and compared the magnitude of BW-evoked ΔF/F (averaged across trials) for each whisker-responsive neuron. In sparse conditions, BW-evoked response magnitude was no different between *Cntnap2^+/+^* and *Cntnap2^-/-^* mice. Noisy conditions reduced the average BW-evoked response magnitude, and *Cntnap2^-/-^*mice showed even smaller BW-evoked ΔF/F than control mice in noisy conditions (Fig. 3g). In sparse conditions, neurons in both *Cntnap2^+/+^* and *Cntnap2^-/-^*mice responded to BW deflection on a similar fraction of trials, but in noisy conditions *Cntnap2^-/-^* neurons responded on fewer trials than control mice (Fig. 3g). Thus, there was no evidence for excess whisker-evoked activity in L2/3 of S1. Instead, we found evidence for increased spontaneous activity (Fig. 3c), which is consistent with the c-fos results, but normal or slightly suppressed whisker-evoked activity, and slightly broader whisker tuning curves.

### Blurred whisker map in *Cntnap2^-/-^* mice

In wild type mice, L2/3 PYR cells tuned to different, nearby whiskers are spatially intermixed in each column, creating local tuning heterogeneity within the whisker map. To test whether whisker map topography is altered in *Cntnap2^-/-^*, we first analyzed this tuning heterogeneity among whisker-responsive PYR cells in one column (Fig. 4a). In sparse conditions, 60% of PYR neurons in each column were tuned to the columnar whisker (CW) in *Cntnap2^+/+^* mice, but only 50% were CW-tuned in *Cntnap2^-/-^* mice (p<0.001, Fisher exact test). When background whisker noise was present, this increase in tuning scatter was even greater (59% CW-tuned in *Cntnap2^+/+^*mice vs. 44% *Cntnap2^-/-^* mice, p<0.001).

**Figure 4.**
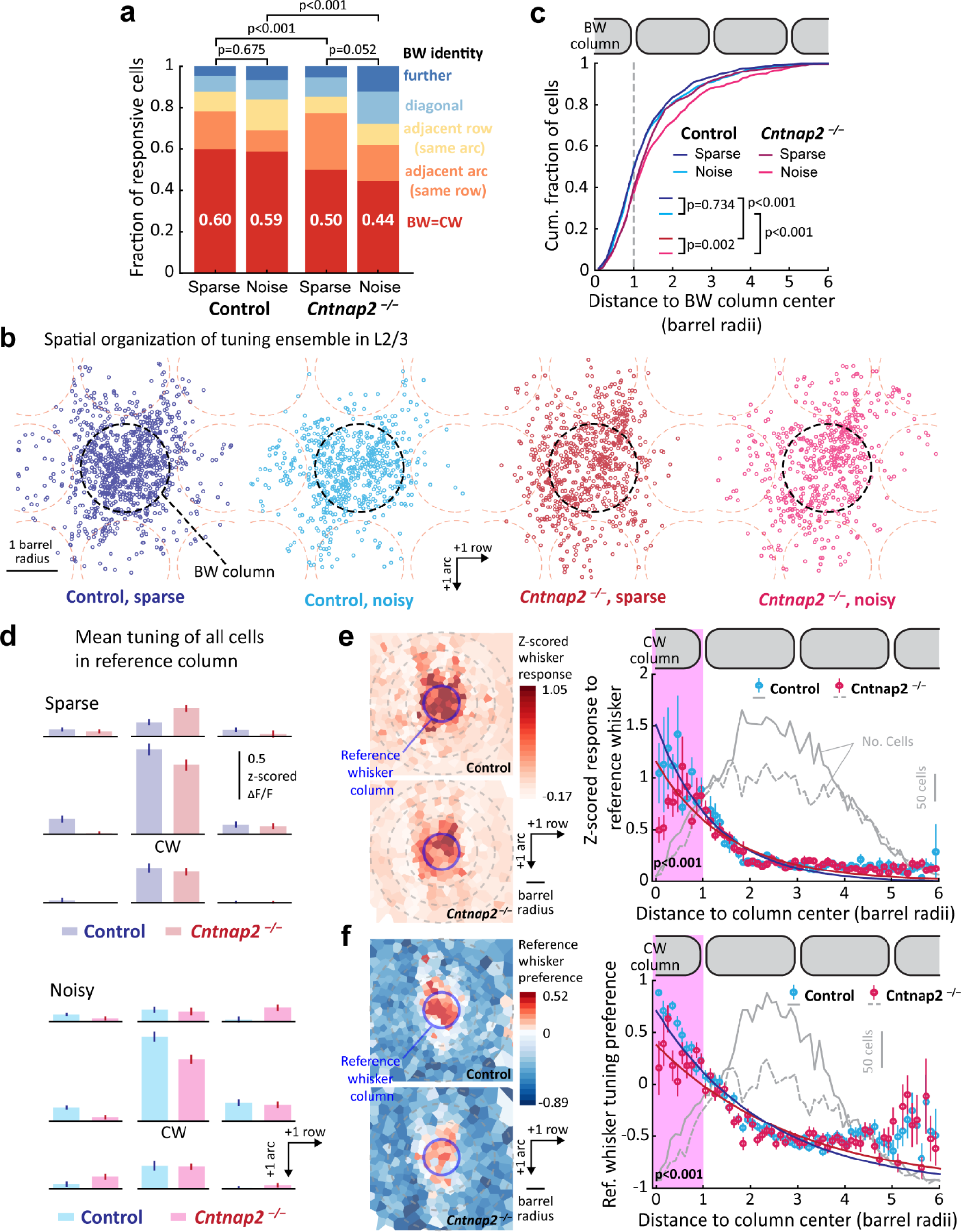
Blurred whisker map in L2/3 of *Cntnap2^-/-^* mice **(a).** BW identity for all responsive cells in a whisker column, by genotype and noise condition. Cell numbers (left to right): 926, 634, 697, 607. Statistics: Fisher’s exact test for BW=CW or not. **(b).** Spatial distribution of the tuning ensemble in control and *Cntnap2^-/-^*mice. Each panel shows the location of each PYR neuron relative to its BW column. Cell numbers (left to right): 1123, 745, 874, 722. Dashed circles show the average location of nearby columns. **(c).** Distance of each responsive cell from its BW column center. Statistics: KS. **(d).** Mean whisker tuning curve for all responsive cells in a column. Whiskers are shown somatotopically, with the columnar whisker (CW) in the center. Responses were z- scored to spontaneous activity in each cell. **(e).** Left: 2D spatial distribution of evoked responses to a given reference whisker, normalized to spontaneous activity in each cell. Cell location is plotted relative to the reference whisker column. Cells were spatially binned using k-means clustering so that each polygonal bin contains 20 cells. Colors show mean CW-evoked response for all cells in the bin. Right: Average response magnitude to the reference whisker for all whisker-responsive neurons, binned by cell distance to the reference whisker column center. This shows the point representation of a whisker among all whisker-responsive cells in L2/3. Navy blue and carmine lines: single-exponential fit to the data. Error bars: SEM. Statistics: rank-sum for data within the whisker column. **(f).** Left: 2D spatial distribution of tuning preference to a reference whisker, plotted as in (e). Color indicates mean CW preference index in each bin. Right: Average preference for a reference whisker among all whisker- responsive neurons, binned by cell distance from the reference whisker column center. This quantifies the tuning gradient across all whisker-responsive neurons. Plotting and statistics as in (e).

Because of the intermixing of tuning in each column, the set of neurons tuned to any given whisker, termed the tuning ensemble for that whisker, is distributed across multiple nearby columns in wild type mice (Clancy et al., 2015; LeMessurier et al., 2019; Wang et al., 2022). To test for differences in this map organization, we analyzed the spatial organization of the tuning ensemble by identifying all cells tuned to a given reference whisker, and plotting the location of these neurons relative to column boundaries defined by the L4 barrel pattern (Fig. 4b-c). In control mice, about half of PYR neurons within a given whisker’s tuning ensemble were located within that whisker’s anatomical column, reflecting the distributed nature of the L2/3 whisker map (sparse conditions: 49.3%; noisy conditions: 49.9%; raw data: Fig. 4b; quantification: Fig. 4c). In *Cntnap2^-/-^* mice, this fell to 39.8% in sparse conditions and 37.4% in noisy conditions (Fig. 4c). This quantification was based on 1123, 745, 874 and 722 cells in control sparse, control noisy, *Cntnap2^-/-^*sparse and *Cntnap2^-/-^* noisy conditions, respectively. This increased dispersion of the tuning ensemble in *Cntnap2^-/-^* was not due to different spatial sampling of imaging fields or neurons between conditions, because subsampling of neurons to ensure an identical columnar distribution of neurons across genotypes and conditions yielded similar results (Fig. 4-Figure Supplement 1a-c). Thus, the whisker map in *Cntnap2^-/-^* mice is even more intermixed and scattered than in control mice.

Consistent with fewer cells tuned for the CW in each column, the mean whisker tuning curve across all whisker-responsive cells in a column was less dominated by the CW in *Cntnap2^-/-^* mice than in control mice, for both sparse and noisy conditions (Fig. 4d). The point representation of a given whisker, defined as the tangential profile of whisker-evoked response magnitude across cortical distance in S1, was blunted in *Cntnap2^-/-^* mice, both in sparse conditions (Fig. 4e) and noisy conditions (Fig. 4-Figure Supplement 2a-b). We also calculated the spatial profile of tuning preference for a given whisker, using a tuning preference index that varies between +1 (cells respond exclusively to that whisker) and -1 (cells respond exclusively to a different whisker). This tuning preference index normally falls off with cortical distance from the whisker’s column center (Wang et al., 2022). This spatial profile was blunted in *Cntnap2^-/-^* mice in sparse conditions (Fig. 4f), but not in noisy conditions (Fig. 4-Figure Supplement 2c-d). Together, these findings show that the L2/3 whisker map in *Cntnap2^-/-^* mice is blurred and weakened on the columnar level, with each column being less sharply tuned for its CW than in control mice.

Background whisker noise did not alter the organization of L2/3 whisker map in *Cntnap2^+/+^*mice, but the presence of noise made the map even more dispersed in *Cntnap2^-/-^*mice.

We also analyzed noise correlations (trial-to-trial covariability) between pairs of co-columnar PYR cells (Kohn et al., 2016). Noise correlations reflect shared spontaneous activity modulation, and are often used to infer shared network connectivity. In wild-type mice, noise correlations drop off with distance between cells in the same column (Wang et al., 2022). *Cntnap2^-/-^* mice showed higher noise correlations than control mice, particularly at 100-150 µm distances (Fig. 4- Figure Supplement 2e). This result may be related to the increased spontaneous activity in *Cntnap2^-/-^*mice, and may also suggest higher local connectivity between co-columnar neurons in L2/3.

### Increased tuning instability in *Cntnap2^-/-^* mice

Sensory tuning can be unstable across days in primary sensory cortex, causing representational drift that may impact sensory computations and perception (Deitch et al., 2021; Driscoll et al., 2022; Schoonover et al., 2021). Some L2/3 pyramidal cells in S1 of wild type mice exhibit unstable whisker somatotopic tuning in expert mice performing well-learned whisker discrimination tasks, and cells tuned to non-columnar whiskers show the highest rates of tuning instability (Wang et al., 2022). Because *Cntnap2^-/-^* mice have more cells tuned for non-CW whiskers (Fig. 4), we hypothesized that *Cntnap2^-/-^* mice may show more unstable whisker tuning.

To test this, we measured tuning stability in task-expert mice, by longitudinally imaging the same PYR neurons across 3-4 sessions spaced 4-7 days apart (Fig. 5a). We compared whisker tuning of single neurons across 1, 2 or 3 session intervals in *Cntnap2^+/+^* mice (Δ1 interval: 6.4 ± 0.2 days; Δ2: 11.9 ± 0.4 days; Δ3: 18.0 ± 0.5 days) and *Cntnap2^-/-^* mice (Δ1: 6.1 ± 0.3 days; Δ2: 12.8 ± 0.4 days; Δ3: 19.0 ± 0.7 days. These experiments were performed in 2 control and 2 *Cntnap2^-/-^* mice in sparse stimulus conditions. Mice maintained high performance across all sessions, and the rate of spontaneous ΔF/F events was stable across sessions, suggesting stable GCaMP8m expression (Fig. 5- Figure Supplement

**Figure 5.**
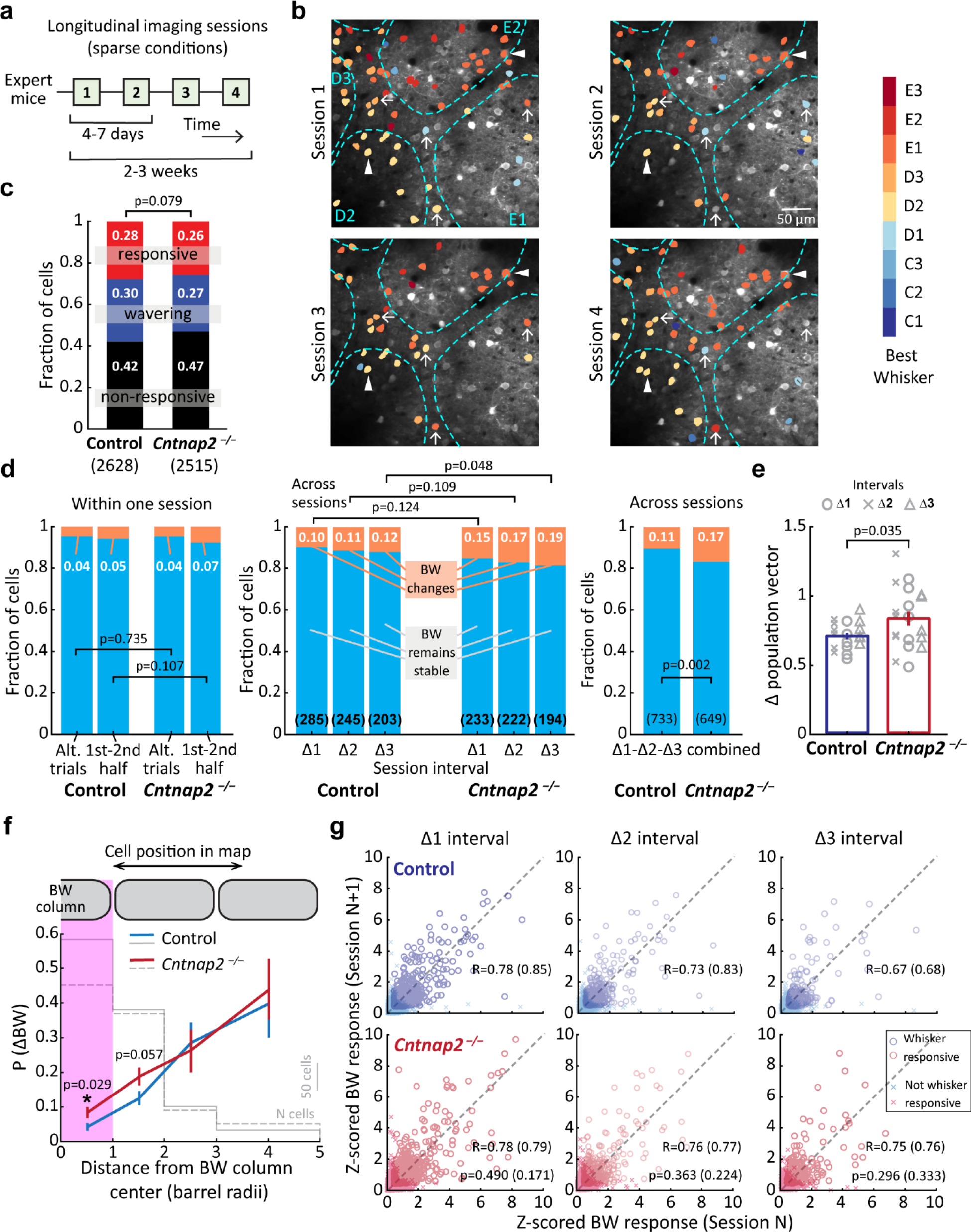
*Cntnap2^-/-^* mice have less stable whisker tuning. **(a).** Timeline of longitudinal imaging sessions from one imaging field. **(b).** Example of 4-session longitudinal imaging from one field, showing barrel boundaries and GCaMP8m-expressing PYR cells color-coded for their BW identity. Arrows: cells changed their BW. Arrowhead: cells kept their BW. **(c).** Changes in responsiveness for cells tracked longitudinally. All intervals were pooled. Numbers are fractions of cells. n: pairs. Statistics: χ^2^. **(d).** The proportion of stably responsive cells whose BW significantly changed within or across sessions. Left: BW changes within the same session. Middle: changes over Δ1, Δ2, or Δ3 session intervals. Right: changes pooled over all intervals. n: pairs. Statistics: Fisher’s exact test for BW changes or not. **(e).** Mean change in whisker-evoked population activity vectors across sessions. See text for explanation. Error bars: SEM. p-values are for genotype factor in unbalanced two-way ANOVA. **(f).** Mean fraction of neurons exhibiting a BW change, as a function of cell distance to its BW column center in session 1. Cells in the magenta area were initially CW-tuned. Data pooled across different intervals. Statistics: rank-sum test between the two behaviors within each bin. Lines with error bars are mean and bootstrapped 95% confidence interval after subsampling to ensure each cell is represented only once. **(g).** Correlation of z-scored BW responses on one session with the previous session, separated by Δ1, Δ2, or Δ3 session intervals. R: Pearson’s correlation coefficient for all cells and responsive cells (in parenthesis). Statistics: permutation test between genotypes.

1a-b). In control mice, 1173 neurons were imaged in at least 2 sessions and 855 neurons in all 4 sessions; in *Cntnap2^-/-^*mice, 1091 neurons were imaged in ≥2 sessions and 811 neurons in all 4 sessions.

Inspection of individual fields revealed many cells that changed whisker tuning, or changed whisker responsiveness, across sessions. Fig. 5b shows an example field from an *Cntnap2^-/-^* mouse, with multiple cells that appear to change BW tuning across sessions (arrows) and others with stable tuning (arrowheads).

To study tuning instability, we first identified cells that were significantly whisker-responsive across multiple sessions. In control mice, across any 2 sessions (Δ1, Δ2 or Δ3 intervals), 42% of neurons were unresponsive in both sessions, 30% wavered between responsive and non-responsive, and 28% were whisker-responsive in both sessions (Fig. 5c). Similar fractions were observed in *Cntnap2^-/-^* mice (Fig. 5c), and for Δ1, Δ2, or Δ3 intervals analyzed individually (Fig. 5- Figure Supplement 1c). We analyzed the stability of whisker tuning for neurons that were responsive in both sessions, by testing for a statistically significant change in identity of the BW (ΔBW), defined as the emergence of a new BW that evoked responses significantly greater than the prior BW by permutation test (with α = 0.05). This approach identifies changes in whisker responses that exceed those expected from trial-to-trial variability and finite trial number. Within a single session, 4-5% of neurons in control mice showed a significant change in BW, as did 4-7% of neurons in *Cntnap2^-/-^* mice (Fig. 5d, left), matching the expected false positive rate of our statistical test. Over Δ1, Δ2, or Δ3 intervals, 10%, 11%, and 12% of cells showed significant BW changes in control mice. Over the same intervals in *Cntnap2^-/-^* mice, 15%, 17% and 19% of cells showed significant BW changes. Pooling Δ1, Δ2, and Δ3 intervals together, *Cntnap2^-/-^* mice showed a significantly higher fraction of cells with ΔBW than control mice (p=0.002, Fisher exact test) (Fig. 5d, right). Thus, whisker tuning was less stable in single PYR cells in *Cntnap2^-/-^*mice.

To test whether these tuning changes degraded the population code in S1 for whisker stimuli, we calculated the population activity vector evoked by deflection of each CW within a given imaging field. This was defined as the vector of mean z-scored ΔF/F for each cell in the field (whether whisker responsive or not), normalized to unit length. We then computed the deviation of these population activity vectors across sessions (defined as Euclidean distance between vector endpoints).

*Cntnap2^-/-^* mice showed more deviation of population vectors across sessions, indicating that population coding of whisker deflection is less stable in *Cntnap2^-/-^* mice (Fig. 5e).

In wild type mice, whisker tuning instability is spatially organized within the L2/3 whisker map, with non-CW tuned neurons having much more unstable tuning than CW-tuned neurons (Wang et al., 2022). We observed this same relationship in control mice in the current data set (Fig. 5f). *Cntnap2^-/-^*mice showed normal rates of tuning instability among non-CW tuned neurons, but significantly increased tuning instability for CW-tuned neurons (Fig. 5f). Increased tuning instability in *Cntnap2^-/-^* mice was not due to broader whisker tuning or weaker BW response magnitude, which are two factors that influence tuning instability in wild-type mice (Fig. 5- Figure Supplement 1d-e) (Wang et al., 2022). Thus, for *Cntnap2^-/-^* mice, excess instability specifically arose from neurons located within their BW column.

Finally, we tested whether S1 neurons in *Cntnap2^-/-^*mice may also show more unstable response magnitude across days, distinct from unstable tuning. This hypothesis is motivated because many ASD genes are involved in activity-dependent plasticity and homeostasis (Mullins et al., 2016; Nelson and Valakh, 2015), and several ASD mouse models show impaired cellular homeostasis in sensory cortex, including *Cntnap2^-/-^* (Blackman et al., 2012; Fernandes et al., 2019; Tatavarty et al., 2020). In wild type mice, L2/3 PYR neurons show remarkably consistent whisker-evoked response magnitude across days (Margolis et al., 2012). To test whether this is altered in *Cntnap2^-/-^* mice, we compared the BW-evoked ΔF/F magnitude for the same cell across sessions at Δ1, Δ2, and Δ3 intervals. BW response magnitude was highly correlated between sessions, both considering all cells in the imaging field or only significantly responsive cells (Fig. 5g). This correlation value was not different between *Cntnap2^-/-^* and control mice for any imaging interval (Fig. 5g), or when Δ1, Δ2, and Δ3 intervals were pooled (Fig. 5- Figure Supplement 5f). Shuffling cell identities between sessions dropped correlation to chance, as expected (Fig. 5- Figure Supplement 5g). Thus, *Cntnap2^-/-^* mice showed normal stability of response magnitude, even though an increased fraction of cells showed instability of whisker somatotopic tuning.

## Discussion

*Cntnap2^-/-^* mouse sensory cortex provides a strong test of the E-I ratio / hyperexcitability model of autism, because these mice exhibit clear PV circuit hypofunction in L2/3 of sensory cortex, including reduced PV cell number, reduced feedforward inhibition, and elevated E-I ratio in S1 (Antoine et al., 2019; Deemyad et al., 2022; Peñagarikano et al., 2011; Vogt et al., 2018). While the simple prediction from the canonical E-I ratio hypothesis is a hyperexcitable cortical network with elevated spike rates, synaptic excitation is also reduced in *Cntnap2^-/-^* mice, which may maintain normal spiking (Antoine et al., 2019). Direct evidence for excess spiking in *Cntnap2^-/-^* sensory cortex has been inconclusive: c-fos measurements and multi-unit recordings have suggested hyperexcitability in S1 and A1 (Balasco et al., 2022; Scott et al., 2022), but single-unit recordings in S1 and V1 have shown normal and reduced sensory-evoked spike rates, respectively (Antoine et al., 2019; Del Rosario et al., 2021). Our findings combine 2-photon calcium imaging and c-fos staining to clearly show that *Cntnap2^-/-^* mice exhibit a slight increase in spontaneous activity, but no increase in sensory-evoked spiking in L2/3 of S1. Instead, *Cntnap2^-/-^*mice exhibit several forms of neural coding degradation, including a blurred somatotopic map, broader whisker tuning, abnormally high noise correlations, and reduced coding stability. These coding phenotypes may generate sensory perceptual impairments in *Cntnap2^-/-^*, without excess spiking.

In our study, c-fos and GCaMP imaging gave consistent results, despite different assay conditions (anesthetized vs. awake), different activity time scales, and different cell type specificity (all L2/3 cells for c-fos, and excitatory L2/3 cells for Ca^2+^ imaging). Elevated spontaneous activity in *Cntnap2^-/-^* mice was evident as higher density of c-fos-(+) neurons in the unstimulated hemisphere in *Cntnap2^-/-^* vs. control mice, and in increased GCaMP8m ΔF/F signals during blank (no-stimulus) epochs. In contrast, c- fos-(+) neuron density in whisker-stimulated columns was not different between *Cntnap2^-/-^*and control, either in absolute terms or normalized to the non-stimulated hemisphere, suggesting that raw magnitude and signal-to-noise ratio of whisker responses were not elevated. This was robustly confirmed in GCaMP8m imaging, which showed no elevation in fraction of whisker-responsive cells or whisker-evoked response magnitude (mean ΔF/F), and even a slight depression in noisy conditions (Fig. 3g-h). These findings are consistent with prior *in vivo* single-unit spike recordings that showed no changes in whisker-evoked spike rate in L2/3 excitatory neurons in S1 (Antoine et al., 2019), and weaker visually-evoked spiking in V1 of *Cntnap2^-/-^* mice, accompanied by reduced detection sensitivity for visual stimuli (Del Rosario et al., 2021).

Thus, S1 and V1 of *Cntnap2^-/-^* mice show PV hypofunction without excess spiking. This combination of circuit phenotypes is a common motif that also occurs in V1 of *Ube3a^m-/p+^* mice, V1 and S1 of *Fmr1^-/-^* mice, and V1 and S1 of *Syngap1^+/-^*mice (Arnett et al., 2014; Goel et al., 2018; Juczewski et al., 2016; Michaelson et al., 2018; Wallace et al., 2017b). In a broad review of ASD mouse models (Monday et al., 2023), PV hypofunction in sensory cortex is only rarely associated with excess spiking, being observed in S1 and V1 of *Shank3b^-/-^* mice (Chen et al., 2020; Pagano et al., 2023) and in A1 and non-whisker S1 of *Fmr1^-/-^*mice (Rotschafer and Razak, 2014; Zhang et al., 2014). Outside of S1, mPFC of *Cntnap2^-/-^* mice also exhibits reduced synaptic inhibition (Lazaro et al., 2019), with normal mean firing rates during locomotion (Lazaro et al., 2019), and higher and more variable spontaneous activity but lower social odor-evoked responses (Levy et al., 2019). These phenotypes strongly resemble our findings in S1. How cortex can maintain mean firing rates despite reduced inhibition may seem puzzling, but an array of homeostatic plasticity mechanisms exist in cortex that stabilize firing rate in this way (Antoine et al., 2019; Gainey and Feldman, 2017). At older ages than studied here, *Cntnap2^-/-^* mice begin to exhibit spontaneous seizures, indicating that excess spiking is generated somewhere in the brain in older animals (Peñagarikano et al., 2011).

*Cntnap2^-/-^* mice show atypical sensory processing for touch, audition, olfaction and vision, including impaired auditory gap detection, impaired novel odor identification, and impaired tactile discrimination of textured objects (Balasco et al., 2022; Gordon et al., 2016; Truong et al., 2015). These behaviors involve sensory discrimination, and could reflect degraded discriminative coding in sensory cortex. We tested for degraded coding because lack of PV inhibition can acutely broaden sensory tuning of pyramidal cells (Aizenberg et al., 2015; Atallah et al., 2012), and because during development, PV circuits regulate critical periods that are essential for experience-dependent refinement of precise excitatory circuits, receptive fields and maps (Hensch, 2005). We observed slightly broader whisker tuning in *Cntnap2^-/-^* mice (Fig. 3e-f), and a substantial blurring of the whisker map in the form of increased intermixing of differently tuned neurons in each S1 column (Fig. 4b-c). As a result, somatotopic tuning for each individual column was broader (Fig. 4f), and the point representation of each whisker was less sharp (Fig. 4e), yielding a L2/3 map with degraded functional columnar organization. Experience in juvenile mice normally sharpens columnar organization in L2/3 of S1

(LeMessurier et al., 2019), suggesting that *Cntnap2^-/-^* mice may have a deficit in experience-dependent refinement of the whisker map.

Abnormal somatosensory maps have been detected in people with autism (Coskun et al., 2009). In ASD mouse models, sensory maps are degraded in *Fmr1 ^-/-^* mice (Antoine et al., 2019; Arnett et al., 2014; He et al., 2019; Juczewski et al., 2016; Rotschafer and Razak, 2013), *Mecp2* overexpression mice (Zhou et al., 2019), and *En2 ^-/-^*mice (Allegra et al., 2014), and abnormally broad single-neuron sensory tuning has been observed in *Fmr1 ^-/-^*, *Mecp2* deletion, and *Ube3a^m-/p+^* mice (Banerjee et al., 2016; Goel et al., 2018; Juczewski et al., 2016; Rotschafer and Razak, 2013; Wallace et al., 2017a), but this is not universal in all ASD models (Garcia-Junco-Clemente et al., 2013; Krishnan et al., 2015; Ortiz-Cruz et al., 2022; Wadle et al., 2023). This degraded neural coding, both on the single-cell and population levels, may contribute to impairments in sensory discrimination in autism (Goel et al., 2018; He et al., 2021; Puts et al., 2014). The increased noise correlations that we observed within each S1 column in *Cntnap2^-/-^*mice (Fig. 4- Figure Supplement 2e) and reduced trial-to-trial reliability (Fig. 3h) are also expected to interfere with accurate sensory decoding.

In our data, weakened response magnitude and blurred whisker map topography were observed in both sparse and noisy sensory conditions, but these effects were stronger in noisy conditions (Fig. 3e-h and Fig. 4a-c). Noisy conditions were intended to test whether external (sensory) noise disrupts neural processing in ASD. One of the prominent sensory features of ASD is the difficulty in detecting speech in background noise (Alcántara et al., 2004; Schelinski and von Kriegstein, 2020). While this is likely to partly reflect a higher level of endogenous neuronal noise (internal noise) in ASD (Rubenstein and Merzenich, 2003; Simmons et al., 2009), abnormalities in filtering out background sensory noise (external noise) may also occur in ASD individuals (Mihaylova et al., 2021; Park et al., 2017), and may reflect increased sensory-evoked spiking and reduced temporal precision of sensory signals at subcortical levels (Nakajima et al., 2019). Our finding that background tactile noise further degrades map topography (Fig. 4b-c), and further weakens BW response magnitudes (Fig. 3g) and trial-to-trial reliability (Fig. 3h) suggest that inadequate management of external noise could be an independent component in the pathogenesis of ASD.

The stability of neural codes has not generally been examined in ASD, but is another factor that may impact sensory and cognitive function. Tuning instability (representational drift) is common in non-topographic, associative cortical areas, but also occurs in S1 and V1 cortex (Deitch et al., 2021; Driscoll et al., 2022; Schoonover et al., 2021; Wang et al., 2022). Sensory tuning drift can interfere with stable decoding of population activity in sensory cortex (Schoonover et al., 2021), although this problem is eliminated if drift is constrained to non-coding dimensions (Deitch et al., 2021). While the function of tuning instability and representational drift is not known, it may relate to plasticity, memory consolidation, or contextual processing (Driscoll et al., 2022; Micou and O’Leary, 2023). In autism, it is possible that lower-than-normal instability indicates overly rigid neural coding, perhaps related to behavioral rigidity or cognitive inflexibility phenotypes, while higher-than-normal instability could impair sensory decoding by higher cortical areas, blurring perception.

We assessed tuning instability across 2-3 weeks in expert mice consistently performing the whisker task, and observed a significant increase in *Cntnap2^-/-^* mice in the fraction of L2/3 pyramidal cells that significantly changed their BW across sessions (Fig. 5d), which increased session-to-session variability in a simple population measure of whisker coding (Fig. 5e). Unlike in wild type mice, where instability occurs almost exclusively for neurons tuned for non-columnar whiskers (Wang et al., 2022), in *Cntnap2^-/-^* mice, excess instability was observed for neurons that were tuned for the CW in the first imaging session, meaning that neural tuning switched back and forth between CW and non-CW whiskers, which rarely occur in control mice (Fig. 5g). Thus, *Cntnap2^-/-^* mice exhibit increased instability in both columnar core and non-columnar surrounds of the whisker tuning ensemble (Wang et al., 2022), which could impair accurate decoding of whisker stimuli. Tuning instability may reflect enhanced instability at the cellular and synaptic level. Accelerated turnover of cortical dendritic spines occurs in several ASD mouse models (Nakai et al., 2018), including in *Cntnap2^-/-^* mice, where spines on the apical dendrite of L5b neuron show enhanced loss over a 4-day period (Gdalyahu et al., 2015).

Together, these findings suggest that tactile sensory deficits in *Cntnap2^-/-^* mice do not result from hyperexcitability and excess spiking in S1, but from degraded and slightly weakened neural coding, including broadened single-unit tuning, a blurred somatotopic map, and increased tuning instability that may impair accurate sensory decoding.

## Materials and Methods

### Animals

Procedures were approved by the UC Berkeley Animal Care and Use Committee, and followed NIH guidelines. *Cntnap2^-/-^* (JAX 017482) and *Cntnap2^+/+^* control (JAX 000664) mice were purchased from the Jackson Laboratory at 2 months of age, and were initially housed in cohorts of three or fewer mice, until craniotomy surgery at 2.5-3 months of age, after which mice were housed singly. Each cage had a running wheel (Mouse Igloo #K3327, Bio-Serv), and mice were maintained on a 12/12 light-dark cycle with humidity 30-70% and temperature 20-26 °C. All behavior training and experiments were conducted during the dark (active) cycle. All experiments were finished before 5 months of age, to avoid the spontaneous seizures that emerge in *Cntnap2^-/-^*animals over 6 months of age (Peñagarikano et al., 2011).

5 control and 5 *Cntnap2^-/-^* mice, of either sex, were used for 2-photon imaging. Of these, 4 of each genotype were imaged in both sparse and noisy conditions, while the remaining mice were imaged only in sparse conditions. 2 control and 2 *Cntnap2^-/-^* mice were used for longitudinal imaging. It was not possible to be blind to mouse genotype, because of behavioral differences between Cntnap2^-/-^ and control mice during acclimation to behavior training.

### Cranial window surgery and viral injection

At 2.5-3 months of age, mice were anesthetized with isoflurane (1-1.5% in O2) and dexamethasone (2 mg/kg), enrofloxican (5 mg/kg), and meloxicam (10 mg/kg) were administered. A stainless-steel head holder with 6 mm aperture was affixed to the skull using cyanoacrylate glue and dental cement. The D1- 3 whisker columns in S1 were localized using transcranial intrinsic signal optical imaging (Drew and Feldman, 2009; Grinvald et al., 1986). A 3 mm diameter craniotomy was made centered on the D2 column. We used two viral strategies to express GCaMP8m in L2/3 PYR neurons. In some mice (2 control and 2 *Cntnap2^-/-^*), we co-injected AAV9-syn-jGCaMP8m-WPRE (Addgene # 162375-AAV9) and AAV1- mDlx-NLS-mRuby2 (Addgene #99130-AAV1) to drive pan-neuronal expression of GCaMP8m and interneuron-specific expression of mRuby2 (Chan et al., 2017). We imaged GCaMP8m in the green channel and mRuby2 in the red channel, and only analyzed mRuby2-negative neurons. In the remaining mice, we co-injected with AAV1-syn-FLEX-jGCaMP8m-WPRE (Addgene # 162378-AAV1) and AAV9.CamKII 0.4.Cre.SV40 (Addgene # 105558-AAV9), which limits GCAMP8m expression to excitatory neurons. AAV injections were made at 250 µm and 350 µm subpial depth, at 3-4 locations in S1 surrounding the D2 column. After AAV injection, a chronic cranial window (3 mm diameter glass coverslip, #1 thickness, CS-3R, Warner Instrument) was attached with dental cement. After surgery, buprenorphine (0.1 mg/kg) was administered for post-operative analgesia.

### Behavioral apparatus and behavioral monitoring

Mice performed the behavioral task daily, 5 days per week. At the start of each behavior session, mice were transiently anesthetized with isoflurane and head-fixed under the 2-photon microscope. 9 whiskers (rows C-E, arcs 1-3) were inserted into a 3 x 3 array of calibrated piezoelectric actuators, centered on the D2 whisker. Whiskers were not trimmed, and were threaded into tubes on the piezos, held by soft glue. Deflections were applied 5 mm from the face. A drink port with capacitive lick sensor recorded licks. Paw guards prevented paw contact with whiskers, piezos, or drink port. After whisker insertion, anesthesia was stopped, and mice recovered from anesthesia and began the behavioral task.

Training was performed in total visual darkness (using 850 nm IR illumination for behavioral monitoring). Uniform white noise (77.4±0.5 dB) was continuously applied to mask sounds from piezo actuators and drink port opening. The task was controlled by an Arduino Mega 2560, which monitored licking, dispensed reward, and governed trial timing, with online user control via custom routines in Igor Pro (WaveMetrics). Mice self-initiated each trial by suppressing licking. A given behavioral session used either sparse stimulus trials, or noisy stimulus trials (see below).

For quantification of behavioral performance, d-prime for detection of S+ stimuli was defined as: d’ = z(hit)-z(false alarm) where z=inverse of the normal cumulative distribution function with mean=0 and standard deviation=1. Reaction time was defined as the time between the onset of whisker cue and the first lick in hit trials.

### Sparse session trial structure

Each trial consisted of a 1 s baseline period, 1.5 s pre-stimulus period, 0.5 s whisker cue stimulus period, and 1.5 s response window. One randomly chosen stimulus was applied per trial: either one of the 9 single whisker cues, the all-whisker deflection, or a blank (no stimulus). Whisker cue stimuli consisted of ramp-return rostrocaudal deflections (300 µm, 5 ms rise/fall time, 10 ms duration), applied in a train of 5 deflections (100 ms inter-pulse interval, 500 ms train duration). Trains were used because they evoked more reliable GCaMP signals in L2/3 neurons than single-deflection stimuli. The all-whisker stimulus consisted of simultaneous whisker cue stimuli delivered across all 9 whiskers.

The response window began at the end of the whisker cue deflection. On S+ trials (all-whisker stimuli), water reward (2-4 µl) was automatically dispensed 300 ms into the response window. Licking was not required to dispense reward. Water was not dispensed on S– trials. Licking above a threshold rate during the response window was defined as a lick response, and scored as a hit on S+ trials and a false alarm (FA) on S– trials. FAs and misses were not rewarded or punished. Each trial was followed by a 2 ± 1 s inter-trial interval (ITI) before the mouse could initiate the next trial. Thus, whisker stimuli were separated by > 6.5 ± 1 s in the sparse condition.

### Noisy session trial structure

For noisy stimulus sessions, trial structure was the same as for sparse stimulus sessions, except that random, small-amplitude, single-whisker deflections (90 µm, which was 30% of whisker cue stimulus amplitude) were applied every 200 ± 100 ms on every trial. These noise stimuli were applied on randomly interleaved whiskers, and started during the prestimulus period and lasted till the end of trial, including the whisker cue stimulus period. No whisker noise was applied during inter-trial intervals or during the baseline period at the start of each trial. All whiskers were randomly interleaved for noise stimuli, independent of which whisker was chosen for the main whisker cue stimulus on that trial.

Noisy sessions were not presented during initial training. Once mice were trained, noisy sessions were presented on a subset of days for Ca^2+^ imaging, intermixed with sparse stimulus sessions.

### Training stages

1-2 weeks after cranial window implantation, mice began water regulation to provide motivation for training. Daily water intake was reduced to an individually determined volume (0.7-1.0 mL) to achieve 85% of ad lib body weight. Mice received water rewards for correct responses during training, and the balance of the water budget was provided after each training session. Mice had free access to food, and weight and health were monitored daily.

Training and imaging were conducted in parallel for each pair of control and *Cntnap2^-/-^*mice.

Training proceeded in stages. In Stage 1 training, mice were acclimated to head-fixation and presence of the water port. *Cntnap2^-/-^*mice took consistently longer to habituate to head fixation, which prevented the experimenter from being blind to genotype. In Stage 2, mice learned to lick for water rewards (2-4 µl) cued by a blue LED mounted on the lick port. In Stage 3, S+/S– training began, using only all-whisker deflection (S+) and blank stimuli (S-). The LED still flashed at the time of water delivery. Over days, mice learned to lick to the S+ stimulus, evidenced by an advance in lick timing from after LED onset to before LED onset, as well as by a reduction in FA licks on S- trials. This training stage continued until FA rate fell below 50%, and >50% of licks on S+ trials occurred prior to the LED cue. In Stage 4, the final full behavioral task was implemented by introducing the other 9 S– stimuli and removing the LED cue. Training was only performed using sparse stimulus conditions.

Imaging sessions began when mice reached stable Stage 4 performance with >900 trials per session, including ∼10-20% S+ trials. Whiskers were not trimmed, and remained intact throughout the experiment.

### Two photon imaging

2-photon imaging took place 4-6 weeks after viral injection. Imaging was performed with a Moveable Objective Microscope (Sutter) and Chameleon Ultra II Ti:Sapphire mode-locked laser (Coherent). GCaMP8m and mRuby2 were excited at 920 µm. Scanning utilized one resonant scanner (RESSCAN-MOM, Sutter) and one galvo scanner (Cambridge Technology). Emission was collected through a 16X immersion objective (0.8 NA, N16XLWD-PF, Nikon), bandpass-filtered with dichroic mirrors (green channel: HQ 575/50, red channel: HQ 610/75, Chroma), and GaAsP photomultiplier tubes (H10770PA-40, Hamamatsu). Laser power at the sample was 30-75 mW. Serial single plane images (512 x 512 pixels, 150-275 µm below dura) were acquired at 7.5 Hz (30 Hz acquisition, 4-frame average) using ScanImage5.6 (Pologruto et al., 2003) (Vidrio). Fields of view of either 305 µm x 305 µm or 406 µm x 406 µm were used.

Each daily session comprised 900-1000 trials. In each mouse, 2-5 different imaging fields were sampled in each type of behavioral session (sparse and noisy). We did not directly compare sparse and noisy conditions in the same field of view. In total, 17 imaging fields were obtained from *Cntnap2^+/+^* mice under sparse conditions, 13 fields from *Cntnap2^+/+^* under noisy conditions, 16 fields from *Cntnap2^-/-^* under sparse conditions, and 11 fields from *Cntnap2^-/-^*under noisy conditions. Longitudinal imaging was performed on 2 *Cntnap2^+/+^*and 2 *Cntnap2^-/-^* mice. For longitudinal imaging, 3-4 fields were imaged in each mouse, only under sparse conditions, with each field re-imaged 3-4 times at 4-7-day intervals.

### Histological localization of 2-photon imaging fields and cells

Cells imaged in 2-photon experiments were localized relative to L4 barrel boundaries using post-hoc histology. A 2-photon z-stack was collected spanning from the L2/3 imaging plane to the pial surface, at the end of each imaging session. After imaging experiments were complete, the brain was removed and fixed in 4% paraformaldehyde, and the cortex was flattened and sectioned (50 µm thickness) parallel to the cortical surface, with individual sections spanning from the surface blood vessels down through L4. These sections were stained for cytochrome oxidase activity or with fluorophore tagged streptavidin (Streptavidin, Alexa Fluor™ 647 conjugate, S21374) (Sumser et al., 2017). Both of these methods reveal L4 barrels. Sections were digitized and barrel boundaries traced from the L4 sections and aligned to the surface vessels. Imaging fields were then aligned to column boundaries using blood vessels as landmarks.

### c-fos quantification

#### Whisker stimulation

3 *Cntnap2^+/+^* mice and 3 *Cntnap2^-/-^* mice were used in this experiment. Mice were lightly anesthetized with 0.5% isoflurane in O2 combined with the sedative chlorprothixene (0.08 mg, i.p.). Mice were then head-fixed and body temperature was maintained at 37°C. Whiskers A1-4, C1-4, E1-4, β and δ on the right side of the face were inserted in a piezoelectric actuator array. These constitute the stimulated whiskers. The B and D row whiskers on this same side were trimmed to prevent accidental movement by piezoelectric actuators. Whisker stimuli (trains of 5 rostrocaudal ramp-return deflections, 100-ms inter-deflection interval, 0.5-sec train duration) were delivered simultaneously to all of the stimulated whiskers, every 5 seconds for 30 minutes. The whiskers on the unstimulated side of the face remained untouched.

#### Immunolabeling of c-fos and quantification of c-fos-positive cells

Mice were sacrificed 60-90 min after whisker stimulation and transcardially perfused with 4% paraformaldehyde. The cortex was flattened and sectioned parallel to the cortical surface. Sections (50 µm thickness) were cut via freezing microtome and stained with anti-c-fos primary antibody (Cell Signaling Technology 2250T or Novus Biologicals NBP2-50037SS), and nuclei were stained with 4′,6- diamidino-2-phenylindole (DAPI). Barrels were labeled with fluorescent streptavidin (Streptavidin, Alexa Fluor™ 647 conjugate, S21374) (Sumser et al., 2017). Images were obtained using a 20X objective (Plan Apo VC 20X DIC N2) on a Nikon spinning disc confocal microscope (Eclipse Ti microscope with Andor DU- 897 camera) with NIS-Elements AR software. Cell counting and analysis were performed with ImageJ (NIH, Bethesda). c-fos positive cells were counted across multiple sections through L2/3, spanning the 250 µm immediately above L4. Images were aligned to column boundaries revealed by streptavidin using blood vessels as landmarks. For quantification, we only counted the cells within column borders (i.e., ignoring cells located above septa). Cell density (c-fos positive cells per mm^3^) was compared between stimulated (A-C-E) and unstimulated (B-D) columns in the stimulated hemisphere, and in these same columns in the non-stimulated hemisphere. For each whisker row A-E, all individual cortical columns within that row were combined to a single point representing that row in Fig 1b-c.

### Calcium imaging analysis

Analysis used the CaImAn (Giovannucci et al., 2019) algorithm and custom Matlab routines unless stated otherwise.

#### Imaging processing and ROI selection

Movies were corrected for slow X-Y motion using NoRMCorre (Pnevmatikakis and Giovannucci, 2017). Substantial Z-axis movement was not observed and not corrected. Neuronal regions-of-interests (ROIs) were defined using CaImAn with default settings. The CaImAn algorithm recognized 80% of visible cells, and remaining cells were manually annotated using CaImAn’s manually_refine_components function based on the average image. ΔF/F traces were extracted by CaImAn, with F0 defined as the 25^th^ percentile of the fluorescence distribution for that ROI. Only ROIs near stimulated whisker columns were analyzed (defined as ≤ 1.25 barrel radii from the centroid of a stimulated whisker column). We manually inspected and removed neurons with their nuclei filled with GCaMP, which indicated overexpression.

Around 5% of neurons were removed from each imaging field. The total number of imaged neurons were: *Cntnap2^+/+^* sparse: 2751; *Cntnap2^+/+^* noisy: 1583; *Cntnap2^-/-^* sparse: 2136; *Cntnap2^-/-^* noisy: 1815. For longitudinal imaging, we also examined the spontaneous event rate through the imaging sessions and found the population spontaneous activity remained constant (Fig. 5- Figure Supplement 1b), suggesting stable GCaMP expression.

#### Whisker-evoked responses and receptive fields

To avoid lick contamination, ΔF/F responses were only analyzed on non-lick trials. Stimulus-evoked ΔF/F was defined as mean ΔF/F (0-1000 ms after stimulus onset) minus mean baseline ΔF/F. To identify significant whisker responses, we used a permutation test for difference in mean ΔF/F for each whisker relative to blank trials. In each iteration of the permutation test, single-trial ΔF/F data were randomly shuffled between whisker S- and blank trials, and the difference in mean response between these shuffled trial sets was calculated. This was repeated 10,000 times to generate a null distribution. A measured whisker response was considered significant if it exceeded the 95^th^ percentile of this null distribution. *p*-values were corrected for multiple comparisons across all S– stimuli with false discovery rate 0.05 (Benjamini-Hochberg procedure) (Benjamini et al., 2001). A cell was considered whisker-responsive if ≥1 whisker induced a significant positive ΔF/F response. A single trial was defined as responsive if stimulus-evoked ΔF/F exceeded the mean plus one standard deviation of blank trials. Negative ΔF/F responses were replaced with zero.

#### Tuning of individual neurons

The best whisker (BW) was defined as the whisker that evoked the largest mean ΔF/F response and was significantly greater than blanks. For a cell to be classified as non-CW-tuned, the non-CW response had to be statistically greater than the CW response. BW tuning sharpness (Fig. 3f) was defined as (RBW - RW)/( RBW + RW), where RBW = mean ΔF/F to BW, and RW = averaged mean ΔF/F for all other whiskers (whiskers that evoked a negative response were considered as zero). Columnar whisker (CW) preference (Fig. 4f and Fig. 4- Figure Supplement 2c-d) was calculated similarly as (RCW - RW)/( RCW + RW), where RCW = ΔF/F to the CW. Rank-ordered tuning curves were calculated by ranking each stimulus from strongest to weakest within each cell (normalizing to the blank) and then averaging ranked tuning curves across cells. This quantifies tuning sharpness around each cell’s BW, independent of somatotopic organization. For rank-ordered tuning curves, only cells whose BW was the center whisker or a center-edge whisker in the piezo array were included. This ensures that the BW plus 5 or 8 immediate adjacent whisker responses were sampled.

#### Normalized anatomical reference frame for spatial analysis across imaging fields

To project cells into a common columnar coordinate system, ROI coordinates were transformed into a polar reference frame. We first drew a vector from the centroid of a reference column to the ROI. The normalized distance from ROI to column center was calculated as (measured distance) / (distance from column center to column edge along this vector). This gives units of barrel column radii. To determine the angular position for each ROI, vectors were drawn connecting the centroid of each surrounding column to the centroid of the reference column. These vectors defined equally spaced 45° angles in reference space, and ROI angle was determined relative to these vectors.

#### Analysis of tuning stability by longitudinal imaging

ROIs were identified independently for each imaging session by CaImAn. ROIs that corresponded to the same neuron across sessions were registered manually based on the average image for each session. 80.6% of imaged neurons could be traced in at least 2 out of 4 sessions. Neurons that could not be traced tended to be close to the imaging field edge and were obscured by image registration, or exhibited very low activity and thus did not appear in average image.

To assess tuning stability across sessions, we tested for a statistically significant change in BW, by testing whether a new whisker evoked a significantly stronger mean ΔF/F than the prior BW, assessed by permutation test.

When neurons were present across all 4 sessions, each neuron could contribute 3 different Δ1 measurements (1^st^→2^nd^, 2^nd^→3^rd^, and 3^rd^→4^th^), 2 different Δ2 measurements (1^st^→3^rd^, 2^nd^→4^th^) and one Δ3 measurement (1^st^→4^th^). To avoid overcounting the same cell in Δ1 and Δ2 measurements, we randomly subsampled a single Δ1 or Δ2 value for each cell, repeated this 1000 times, and reported mean and 95% confidence interval for these measurements (error bars or shadings in Fig. 5f and Fig. 5- Figure Supplement 1d-e).

#### Population activity vector

To evaluate the stability of single whisker representation in L2/3 across sessions, we calculated the population activity vector elicited by deflection of a single whisker in each session. For a given whisker whose column was present in the imaging field, the mean responses of each cell in the imaging field (N cells) to that whisker were concatenated as a Nx1 vector, and normalized by the L2 norm of the vector. The population activity vector was also calculated in the next imaging session, using the same cells. The Euclidean distance between the two vectors (Δ population vector) was used to quantify the change of single whisker representation in that cell population across sessions. If an imaging field contained more than one whisker column, the Δ population vector was determined by averaging the Δ population vectors for each whisker that was present.

#### Spatial subsampling of ROIs

We validated the whisker map differences between genotypes by performing additional analysis to correct for modest differences in spatial distribution of imaged neurons. To do so, we subsampled the data to generate spatially identical sampling in both genotypes. In each iteration of subsampling, cells were randomly chosen from *Cntnap2^+/+^* or *Cntnap2^-/-^*mice so that the numbers of cells within each whisker column were the same. Data analysis was done for these subsampled cells. We performed 1000 iterations of this subsampling. The resulting mean and 95% confidence intervals are reported in Fig. 4- Figure Supplement 1a-b.

### Statistics

Statistical methods are described in Figure Legends and above. Sample size was not pre- determined. All tests were two-tailed except for permutation tests. Single neurons were the unit N, except as follows: Mouse behavior was quantified by mouse and by behavioral session (Fig. 2c-e, and Fig. 5- Figure Supplement 1a). The fraction of responsive neurons per imaging field, spontaneous activity in longitudinal imaging, and population vectors were analyzed by imaging field (Fig. 3d, 5e, and Fig. 5- Figure Supplement 1b). c-fos quantification was quantified by row and side (Fig. 1c-d).

In violin plots, circle is median, horizontal line is mean, thick vertical line is interquartile range, and thin vertical line is 1.5x interquartile range (Fig. 3f).

Abbreviations: KS, Kolmogorov-Smirnoff test. χ^2^, chi-squared test. Rank-sum, Wilcoxon rank-sum test. Data are presented as mean ± standard error of mean (SEM), except where noted.

### Data Availability

Source data for the figures are provided with this paper. Processed imaging data for this study are available on the Feldman lab GitHub repository.

### Code Availability

Matlab analysis code used for imaging data analysis are available on the Feldman lab GitHub repository.

## Acknowledgments

This work was supported by R37 NS092367 from NIH and SFARI Investigator Award.

## Author Contributions Statement

H.C.W .and D.E.F. designed the study. H.C.W. performed the experiments and analyzed the data. H.C.W. and D.E.F. wrote the manuscript.

---

**Figure 3.– Figure Supplement 1.**
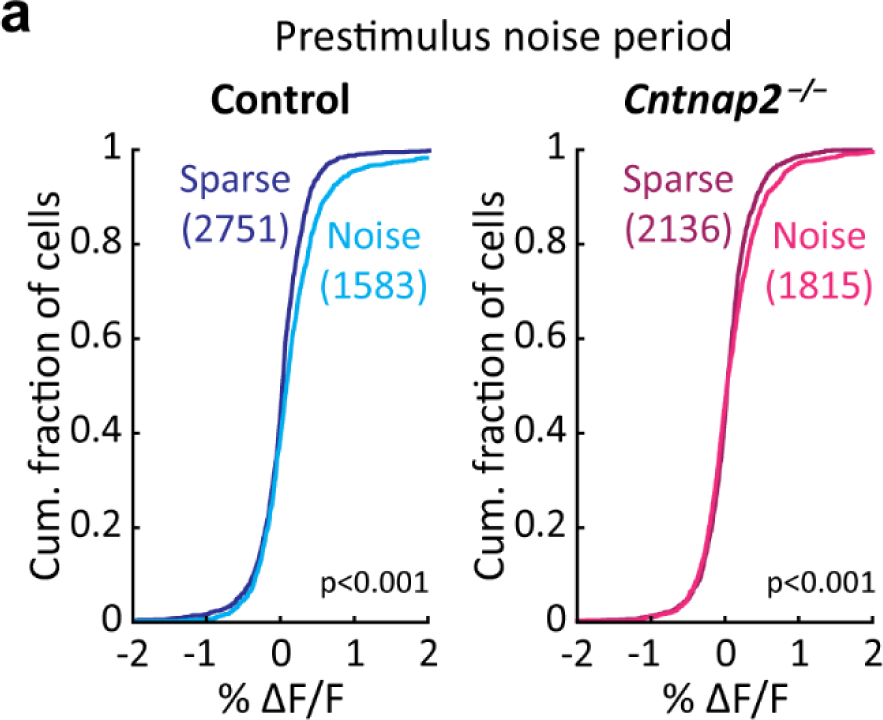
Noisy conditions increase L2/3 PYR activity in the prestimulus period. **(a).** Mean ΔF/F magnitude during the prestimulus period for each cell, by genotype and noise condition. Statistics: KS.

**Figure 4.–Figure Supplement 1.**
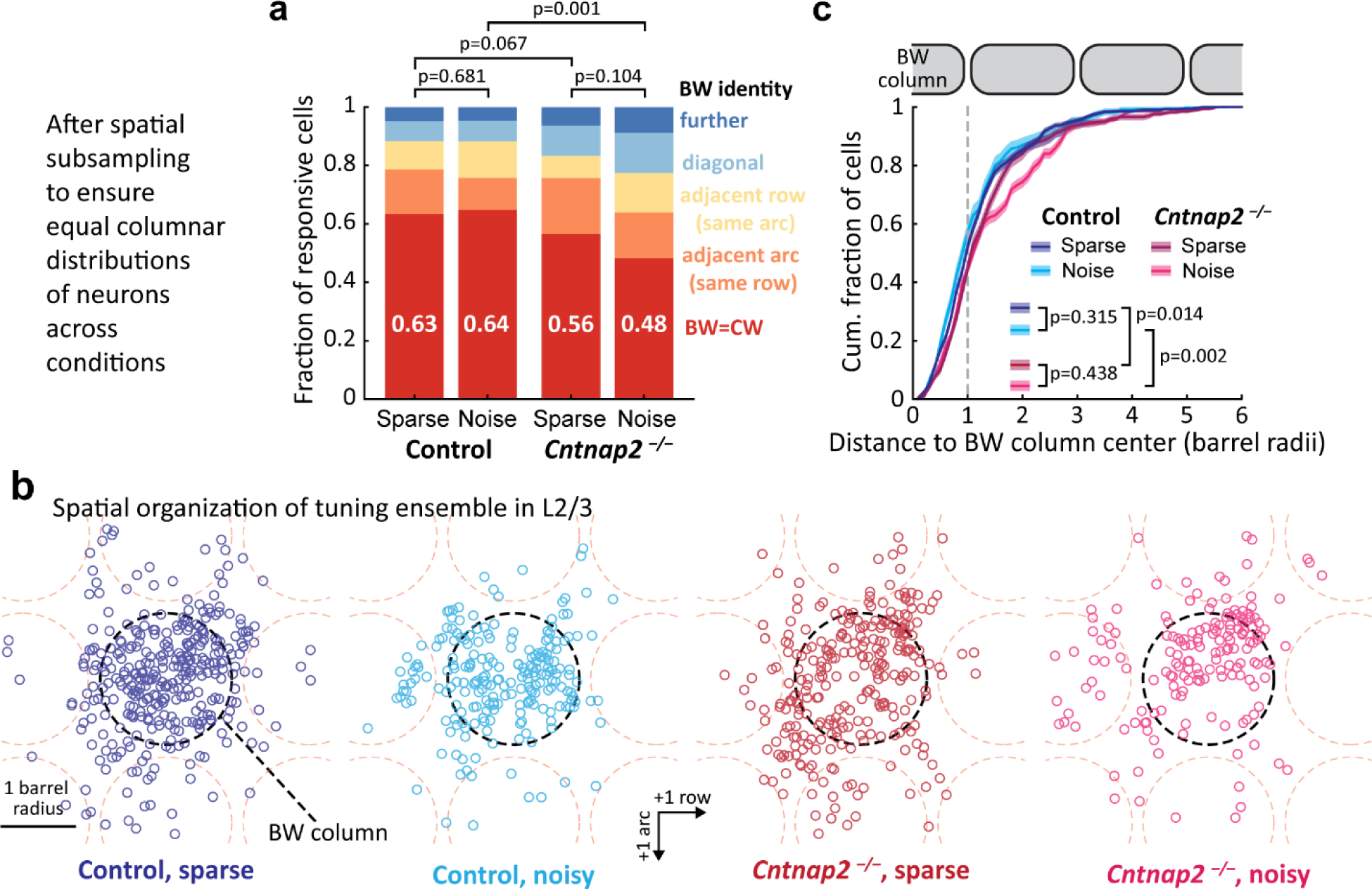
Blurred whisker map after subsampling of neurons to ensure equivalent columnar distribution of neurons across conditions. **(a).** Identity of BW for all responsive cells in a column after spatial subsampling to achieve exactly matched columnar distribution of neurons across genotypes and conditions. Only cells within column boundaries were considered (i.e., septa-related cells were excluded). See Materials and Methods. **(b)**. Spatial distribution of the tuning ensemble in each condition, after the same subsampling procedure. All whisker-responsive cells were included, including column- and septa-related cells. **(c).** Distance of each responsive PYR neuron to its BW column center, after spatial sub-sampling to achieve exactly matched columnar distribution. Shadings show 95% confidence interval from sub-sampling. Same cell population as in (b).

**Figure 4.–Figure Supplement 2.**
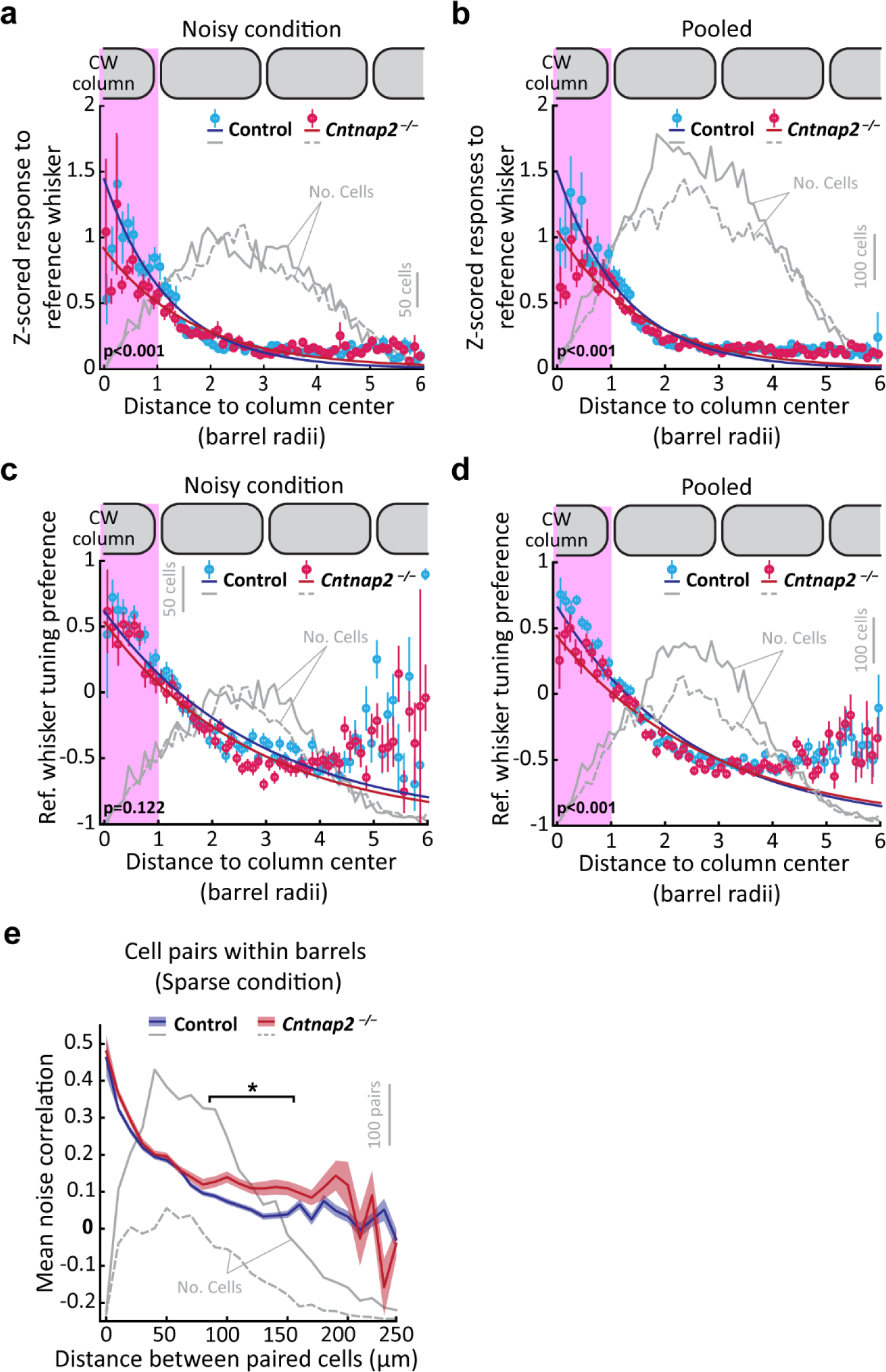
Further quantification of blurred whisker map under noisy conditions. (a-b). Average response magnitude to a given whisker, for all whisker-responsive neurons, as a function of cell distance from that whisker’s column center, in noisy conditions (a) and combining both noise conditions (b). Plotting conventions as in Fig. 4e. Error bars: SEM. Statistics: rank-sum for data within the whisker column. **(c-d).** Average preference for a reference whisker among all whisker-responsive neurons, binned by cell distance from the reference whisker column center, in noisy conditions (c) and combining both noise and sparse conditions (d). Plotting conventions as in Fig. 4f. **(e).** Mean noise correlation for pairs of co-columnar L2/3 PYR neurons, averaged within each distance bin. Asterisk, significant differences between genotypes, by permutation test within each bin. Shading: SEM.

**Figure 5.–Figure Supplement 1.**
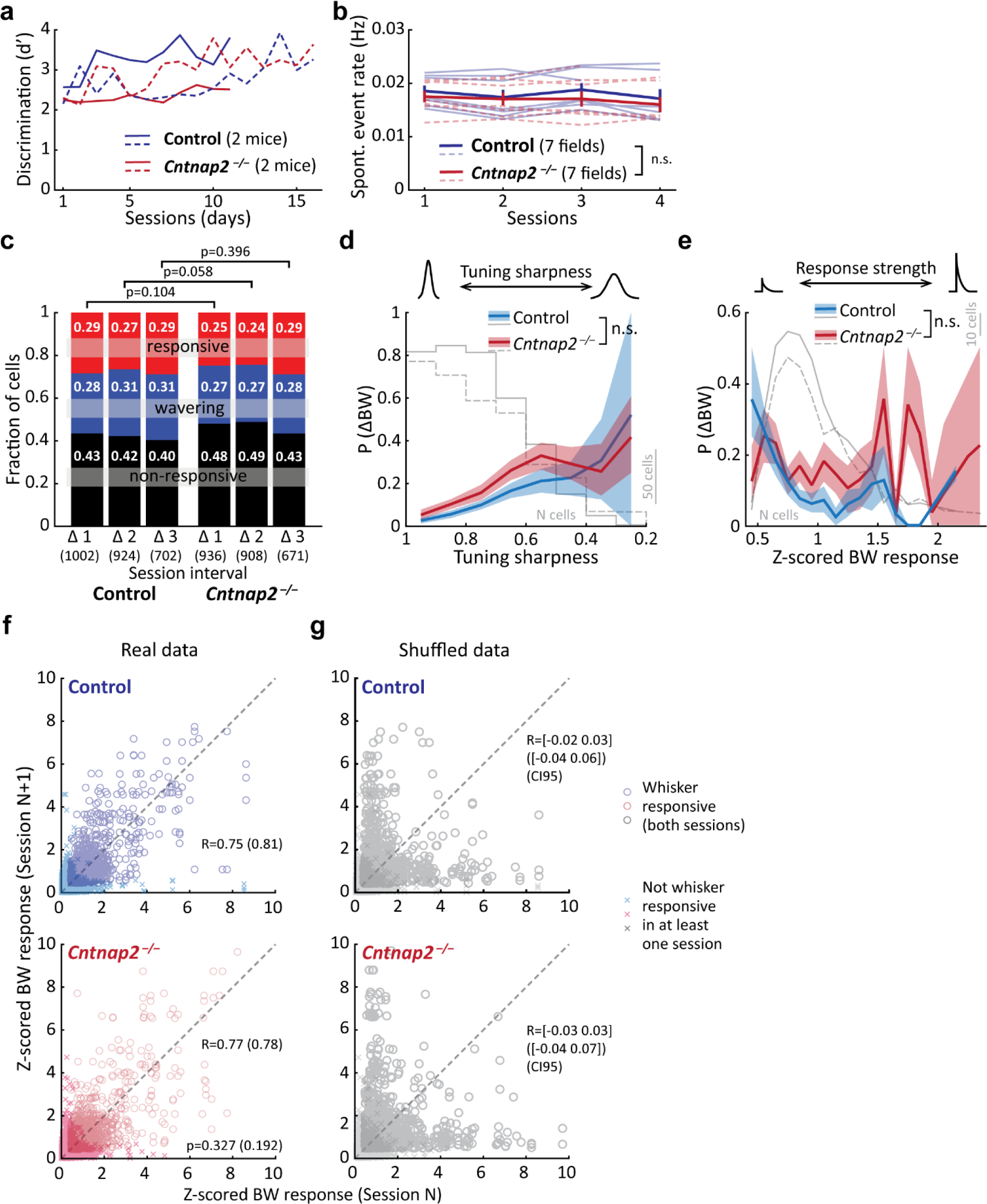
Additional quantification of tuning instability across time. **(a).** Behavioral performance over the period of longitudinal imaging. Each line is one mouse. **(b).** Mean spontaneous event rate for each imaging field across sessions. Each dashed line is one imaging field. Thick lines: grand average ± SEM across fields. **(c).** Changes in responsiveness for cells over Δ1, Δ2, or Δ3 session intervals. Numbers are fractions of cells. n: pairs. Statistics: χ^2^. **(d).** The probability of BW change as a function of initial tuning sharpness. Data pooled across different intervals. Statistics: rank-sum test between the two genotypes within each bin. **(e).** The probability of BW changes as a function of initial BW response magnitude. Data pooled across different intervals. Statistics: rank-sum test between the two genotypes within each bin. In (d) and (e), lines show mean, shading shows bootstrapped 95% confidence interval after subsampling to ensure each cell is represented only once. **(f).** Correlation of z-scored BW responses on one session with the next session, including all pairs of intervals. R: Pearson’s correlation coefficient for all cells, and only for responsive cells (in parenthesis). **(g).** Same analysis as (f), but after shuffling data between the two sessions. R: Confidence interval of Pearson’s correlation coefficient in the shuffled dataset for all cells, and only for responsive cells (in parenthesis).

